# Zika virus infection in a cell culture model reflects the transcriptomic signatures in patients

**DOI:** 10.1101/2024.05.25.595842

**Authors:** Gillian Berglund, Claudia D. Lennon, Pheonah Badu, J. Andrew Berglund, Cara T. Pager

**Affiliations:** RNA Institute, College of Arts and Sciences, University at Albany-SUNY, Albany, NY 12222, USA; Department of Biological Sciences, College of Arts and Sciences, University at Albany-SUNY, Albany, NY 12222, USA

**Keywords:** Zika virus, Transcriptome, Alternative Splicing, Cell Culture, Patients

## Abstract

Zika virus (ZIKV), a re-emerging flavivirus, is associated with devasting developmental and neurological disease outcomes particularly in infants infected *in utero*. Towards understanding the molecular underpinnings of the unique ZIKV disease pathologies, numerous transcriptome-wide studies have been undertaken. Notably, these studies have overlooked the assimilation of RNA-seq analysis from ZIKV-infected patients with cell culture model systems. In this study we find that ZIKV-infection of human lung adenocarcinoma A549 cells, mirrored both the transcriptional and alternative splicing profiles from previously published RNA-seq data of peripheral blood mononuclear cells collected from pediatric patients during early acute, late acute, and convalescent phases of ZIKV infection. Our analyses show that ZIKV infection in cultured cells correlates with transcriptional changes in patients, while the overlap in alternative splicing profiles was not as extensive. Overall, our data indicate that cell culture model systems support dissection of select molecular changes detected in patients and establishes the groundwork for future studies elucidating the biological implications of alternative splicing during ZIKV infection.

## Introduction

Zika virus (ZIKV) is an arbovirus virus belonging to the *Flaviviridae* family *and* Flavivirus genus [1,2]. The virus can cause asymptomatic infections or infections with mild symptoms such as fever, rash, and arthralgia [3–10]. For decades, sporadic ZIKV infections were confined to Africa and Asia, and because of this and the less severe disease symptoms, the virus as a significant human pathogen was overlooked. However, this changed when a ZIKV outbreak occurred in the Yap Island of Micronesia for the first time in 2007 [11] and in the Islands of French Polynesia a few years later infecting about 73% of the inhabitants [12–14]. ZIKV continued to spread rapidly and in 2015 and 2016, the virus caused an epidemic within the South and Central regions of the Americas [1]. This recent epidemic had the highest infection incidence ever recorded and was also associated with unprecedented neurological complications [15,16]. These included Guillain-Barré syndrome, an autoimmune condition which caused paralysis in adults [17–22], as well as congenital malformations such as microcephaly in newborns whose mothers were infected during pregnancy [15,16,23–27]. Besides the severe disease outcomes, the recent epidemic further revealed ZIKV could be spread sexually, via blood and through intra-uterine transmission [28–30]. Although the scale and magnitude of ZIKV infections has declined since 2017, the virus remains an imminent threat and this is particularly concerning given the rapid global expansion of the *Aedes* vector [31], and the lack of approved antiviral drugs and vaccines. Together this underscores the need for more comprehensive studies on the molecular biology of ZIKV infection and the consequence of virus-host interactions.

When ZIKV is injected into the skin during a mosquito blood meal, the virus attaches to cell surface receptors and adhesion factors such as AXL, DC-SIGN, Tyro-3 and in some cases TAM-1 to initiate viral entry through clathrin-mediated endocytosis and membrane fusion [32,33]. Therefore, these receptors define which cell types are permissive to ZIKV, and accordingly, recent studies have described fibroblast, epithelial cells, immature dendritic cells, brain cells, stem cells and blood cells as disease relevant cell types [32,34]. *In vitro* evidence and *in utero* transmission of ZIKV from mother-to-fetus also implicate trophoblasts and macrophages in ZIKV trophism [35]. Additionally, ZIKV has been found to persist in different body fluids including semen, saliva, tears, urine and target organs like the female reproductive tract and immune-privileged sites e.g., eyes, brain, and testes [36–42]. This demonstrates thatZIKV is pan-tropic, with a broad range of target cells.

Following infection, ZIKV infected individuals usually show two phases of disease; acute and convalescent, and the acute phase can further be classified into early and late stages [43]. The acute phase represents days 1 to 6 after the start of disease. In this phase, the virus actively replicates, causes viremia, and activates an initial immune response mediated by innate immunity effectors and IgM [44]. The convalescence phase spans the period from the second week up to about 6 months after disease onset. This stage is marked by high viremia and a more specific and robust interferon (IFN), inflammatory and IgG response [44,45]. For any given period during ZIKV infection, the specific viral processes that takes place and the associated host responses drive distinct transcriptional changes. Such changes are emphasized in a recent study examining ZIKV-infected serum samples from DENV-naive and DENV-immune pediatric patients. Despite prior DENV immunity, the study revealed distinct temporal patterns of gene expression, cell profiles, and inflammatory signatures at the early-acute, late-acute, and convalescent phases (47). These phase-specific molecular signatures, as highlighted by Michlmayr et al., provide broader insights into pathways and biomarkers that could influence therapeutic and diagnostic approaches. However, such detailed information is often inadequately captured in cell culture models [46].

Transcriptomic studies in different clinically relevant cell culture models of ZIKV infection are available [47–52]. They include infections in neuronal and non-neuronal models including neuroblastoma cells, placental trophoblasts, immune cells, and human retinal pigment epithelia (RPE) cells [35,45,53,54]. These analyses give basic understanding of how the gene expression landscape is reprogramed during infection and reveal discrete transcriptomic alterations influenced by cell type, strain of virus and duration of the infection. However, these studies fail to correlate the gene expression changes in *in vitro* cell-based models with a specific clinical infection phase and as a result limit the translational application of these transcriptomic analyses. For example, in human-induced neuroprogenitor stem cells (hiNPCs) infected with ZIKV Brazil (a contemporary isolate) versus ZIKV Cambodia (an ancestral Asian isolate), differential gene expression analysis revealed that both strains strongly modulated immune and inflammatory responses, cell death and growth-related pathways after 48 hours of infection [55]. Likewise, we previously examined transcriptional profiles in a neuronal cell line (SH-SY5Y) following infection with ZIKV MR766 (original ZIKV isolate), ZIKV PRVABC59 (a contemporary strain isolated in Puerto Rico) [56] and DENV serotype 2 (DENV2) (a closely related flavivirus) [47]. From our RNA-seq analysis, we found that 24 hours post-infection, the contemporary ZIKV isolate dramatically influenced the transcriptional landscape compared to the original ZIKV African isolate or DENV2 albeit all three viruses infected cells to the same degree. Notably, ZIKV PRVABC59 upregulated more genes involved in the cellular response while ZIKV MR766 affected genes associated with cellular localization [47].

Besides transcription, the diversity and repertoire of genes in the cell is further expanded through alternative splicing [57]. During viral infection, the cellular splicing machinery may be leveraged for processing viral RNAs and in the process, host splicing, and transcriptional complexity is altered [58,59]. Such alterations in RNA processing have been reported for several viruses and shown to affect genes important for host-virus interactions [59–62]. We previously showed that ZIKV infection could similarly elicit alternative splicing changes in a human neuronal cell line (48). ZIKV infection affected skipped exons (SE), alternative 5’ splice site selection, alternative 3’ splice site selection, mutually exclusive exons and retained intron alternative splicing events with SE events being the most prevalent type of alternative splicing dysregulated by ZIKV infection (48). Although the significance of specific alternatively splicing of cellular transcripts caused by ZIKV infection have not been elucidated, it is likely that mis-spliced mRNAs could impact mRNA stability, localization or translation which could have direct consequences on cellular events contributing to ZIKV-linked pathologies [47,52].

While many studies have previously characterized transcriptomic changes following infection of ZIKV in various cell lines [47,53–55,63–65], few studies have transcriptomic data from human patients and fewer have compared immune and transcriptomic responses between patients and cell models. Although there are a few studies on alternative-splicing changes in ZIKV infected cells [47,52,66], there are no studies on how such changes compare with patient transcriptomic data from the distinct phases of infection. Michlmayr and colleagues [46] collected peripheral blood mononuclear cells (PBMCs) from pediatric patients infected with ZIKV and conducted analyses via RNA-sequencing (RNA-seq), cytometry by time-of-flight (CyTOF), and Luminex cytokine/chemokine multiplex bead array [67]. In this study, we used this publicly available RNA-seq dataset, and further analyzed the dataset for differential gene expression and alternative splicing. We also compared these findings to a cell model of ZIKV infection in A549 human lung adenocarcinoma cells which are highly permissive for ZIKV infections and elicit a robust antiviral response making these cells an acceptable cell culture model in the ZIKV field [68,69].

## Materials and Methods

### A549 cells

Human lung adenocarcinoma cells (ATCC CCL-185) were maintained in Dulbecco’s minimal essential media (Gibco) supplemented with 10% fetal bovine serum (Seradigm, Avantar), 10 mM nonessential amino acids (Gibco), 2 mM L-glutamine (Gibco) and grown at 37°C with 5% CO_2_.

### Zika virus infections

A549 cells were seeded a day before infection and incubated at 37°C overnight. The next day, cells were approximately 80% confluent, and one culture plate was trypsinized and counted to calculate the viral volume needed for a multiplicity of infection (moi) of 10 plaque forming unit (PFU)/cell. Cells were incubated with ZIKV PRVABC59 or PBS for mock infection at 37°C for 1 hour, rocking every 15 minutes. Afterwards, 9 mL of media was added to cells and the ZIKV and mock infected cells were incubated at 37°C. At 48 hours post treatment, infection was validated by plaque assays, and western blot and RT-qPCR of ZIKV protein and RNA levels.

### RNA-seq of mock and ZIKV infected A549 cells

At 48-hours post infection, media from the cells was removed, and cells were washed with cold PBS. Hereafter, a cell lifter was used to remove cells from the dish, which were pelleted briefly at high-speed in a microfuge at room temperature. The PBS supernatant was aspirated, and cells were lysed in 1 mL TRIzol (Ambion by Life Technologies). Total cellular and ZIKV RNA was extracted per the manufacturer’s instructions. DNA remaining in the RNA samples was removed by DNase 1 (NEB) digestion, and then the RNA was ethanol precipitated at -20°C overnight. ZIKV infection was validated by RT-qPCR prior to submitting 1 μg of RNA from three independent experiments to Genewiz (from Azenta) for RNA-seq library preparation and analysis.

### RNA-seq analysis

RNA-seq raw datasets for DENV naïve patients were acquired and downloaded from the NCBI Sequence Read Archive (SRA) Explorer. FASTQ files of technical replicates were merged, and the quality checked with FASTQC (version 0.11.9). Files were then trimmed with fastp (version 0.23.2) using default parameters. Trimmed FASTQ files were aligned to the latest human genome (GRCh38) using STAR (version 2.7.10a) [70]. Raw RNA sequencing datasets from mock and ZIKV infected A549 cells were similarly processed. Figures for differential gene expression and alternative splicing were generated using R (version 4.3.1) and the ggbiplot package in RStudio (2023.06.16; 4.3.1).

### Differential gene expression (DGE)

Differential gene expression was performed in RStudio (2023.06.16; 4.3.1) using DESeq2 (version 1.40.2) [71] and EdgeR (version 4.0.5) [72]. Counts were normalized by variance stabilizing transformation (vst) prior to differential expression. Genes that passed a threshold of padj < 0.05 and log_2_ Fold Change (FC) >|1| were considered significantly differentially expressed. Gene ontology of differentially expressed genes was performed using Panther [73], and heatmaps were produced using the pheatmap package in R.

### Alternative splicing

Replicate multivariate analysis of transcript splicing (rMATS version 4.1.2) [74] was used to study differential alternative splicing. Significant SE events were filtered using the cutoffs of delta Percent Spliced In (ΔPSI) > |0.1| and an FDR < 0.05. Venny (version 2.1.0) [75] was used to find overlapping significant SE events across the Michlmayr *et al*. study [46] and the comparison between the patient samples and the A549 cell line. Gene ontology analysis of significant SE events was performed using Panther (77). All other alternative splicing figures were generated using GraphPad Prism 10 and/or Adobe Illustrator 2024.

### RT-qPCR analysis

To validate the differential gene expression changes revealed by RNA-seq analysis, we first isolated total RNA from mock and ZIKV infected cells with TRIzol reagent (Ambion by Life Technologies) and the RNA Clean and Concentrator kit (Zymo Research). The RNA was DNase-treated using the TURBO DNA-free^TM^ kit (Invitrogen) and reverse transcribed into cDNA with the High-Capacity cDNA Reverse Transcription reagents (Applied Biosystems). For each reaction, 1 μg of RNA sample in 10 μL was used. The cDNA was then used as a template for target gene amplification by qPCR using iTaq Universal SYBR Green Supermix reagent (Biorad) and CFX384 Touch Real-Time PCR system (Biorad). The fold change in RNA expression of an indicated target gene was determined relative to *ACTB* (β-actin mRNA) and was determined from three independent experiments, which were also independent from the RNA-seq of uninfected and ZIKV infected A549 cells. Error bars on RT-qPCR represent ± standard deviation (SD), and statistical significance was determined using an unpaired student t-test. Primer sequences are listed in Table S1.

### Alternative splicing (AS) PCR analysis

For cDNA synthesis, 1μg of RNA extracted from mock and ZIKV PRVABC59 infected cells and DNase treated, as described above, was used. The cDNA was used as template in Taq Polymerase PCR reactions for 34 cycles using High Fidelity Hot Start Taq Polymerase 2x master mix (Thermo Fisher Scientific) and gene specific primers designed to bind regions flanking the included/excluded exon (Table S2). The PCR amplicons were subsequently separated by Fragment Analyzer capillary electrophoresis using the DNF905 1-500 base pair kit (Agilent). Bands were quantified and PSI was calculated as follows: PSI = ((RFU of Inclusion Band)/ (RFU Inclusion Band+RFU Exclusion Band)) × 100%, where RFU represents Relative Fluorescence Units. Graphs for alternative splicing events were made using the GraphPad Prism 9.4.1 software (GraphPad, La Jolla, CA, USA) based on PSI analyses from three independent experiments. Error bars represent ± SD and the statistical significance was determined by unpaired student t test in the program.

### Data availability

The patient data obtained from Michlmayr *et al*. [46] is in Gene Expression Omnibus at GSE129882. The cell model data from this study of uninfected ZIKV PRVABC59 infected A549 human lung adenocarcinoma cells has been deposited at GSE265922.

## Results

### PBMCs isolated from ZIKV infected patients show differential transcriptomic profiles during early and late acute stages of infections

Michlmayr *et al*., collected blood samples from 89 pediatric patients at three timepoints of ZIKV infection namely at early acute (1-3-days after symptom onset), late acute (4-6-days after symptom onset), and convalescent phases (14-21-days after symptom onset) [46]. The participants were aged 2-14 years old and were part of the Pediatric Dengue Cohort Study (PDCS) based in Nicaragua. Patients included in the Michlmayr *et al*., study presented with ZIKV infection between July and August 2016 and were confirmed positive by RT-PCR. Out of the total sample set, 46 of the patients had no prior documentation of DENV infection and tested negative for DENV antibodies. We conducted analyses on 45 of the DENV negative patients that had samples from all three timepoints. The average age for these participants was 8.9 years old and there were 22 males and 24 females [46]. We obtained the publicly available RNA-seq data prepared by Michlmayr and colleagues from Gene Expression Omnibus. We independently processed and mapped the transcripts to the human genome prior to downstream analyses.

In our analyses of differential gene expression across ZIKV infection timepoints, we also observed a temporal pattern that the previous study reported on with each timepoint characterized by a subset of unique genes. To further visualize differences between samples of each infection phase, we performed principal component analysis (PCA) on normalized counts (**Figure 1A**). We observed that samples from late acute infections and convalescent samples overlapped, and the early acute samples were partially separated (**Figure 1A**). We also examined if patient sex influenced the distribution of the samples within each period of the ZIKV infection. Here the samples were further separated by gender, and the PCA plot showed distinct differences in gene expression between females and males (**Figure 1A**). However, after performing separate analyses of differential gene expression and comparing sex differences, we did not find any significant differences in the response to ZIKV infection based on sex. Thus, for downstream analyses both sexes were grouped together.

**Figure 1.**
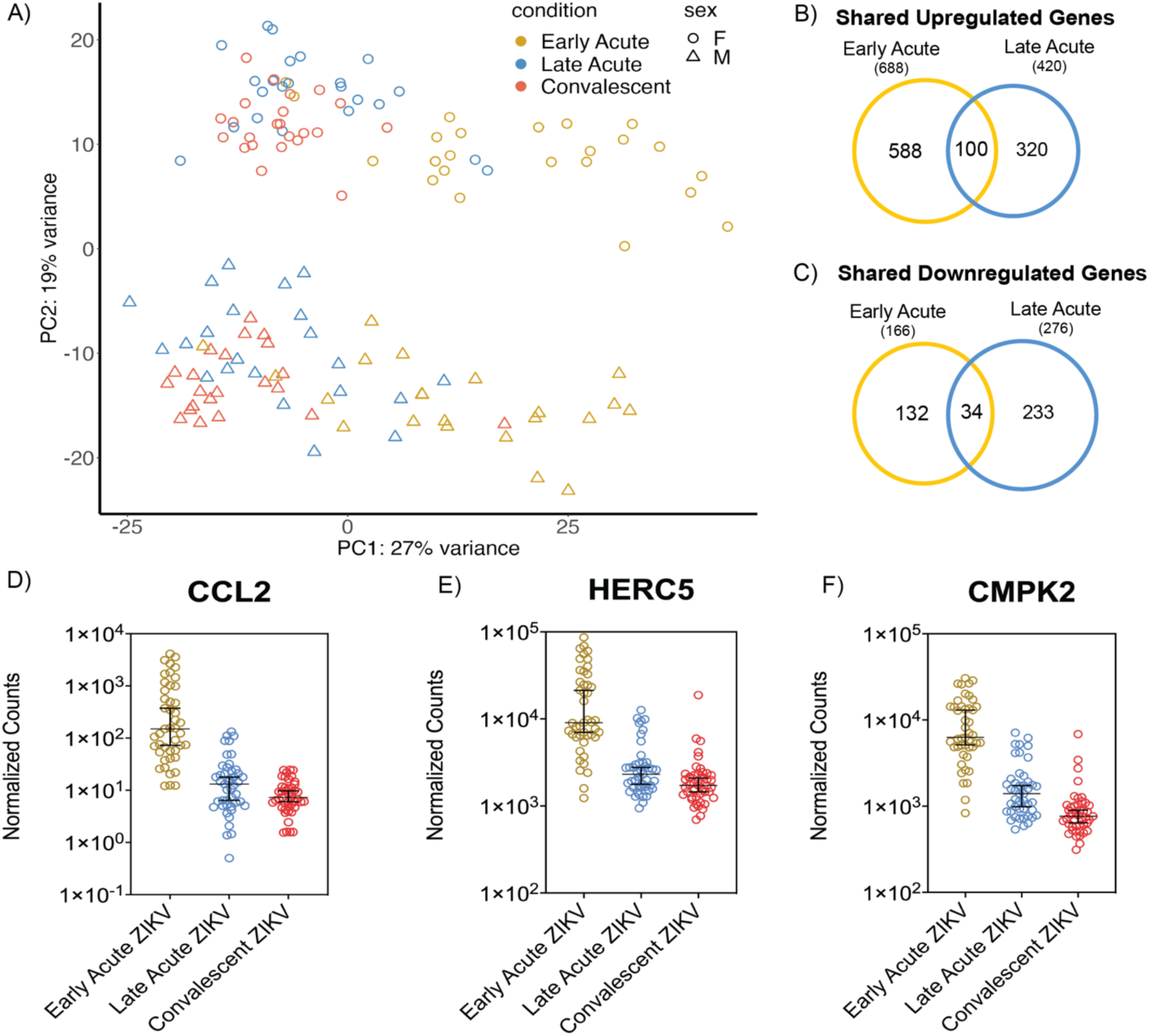
Transcriptome analysis of ZIKV infection at early acute and late acute times in patients. **(A)** Principal component analysis (PCA) of normalized counts. The principal components are colored by infection time (gold – Early Acute, blue – Late Acute, and red – Convalescent) and the reported sex of the patients is represented as circles for female and triangles for male. Venn diagrams show differentially expressed genes at either early or late acute ZIKV infection times relative to the convalescent stage that were either **(B)** upregulated or **(C)** downregulated. The differentially expressed genes are defined as Log_2_ FC > 1. **(D)** Normalized counts of *CCL2* expression across each ZIKV infection stage. The average Log_2_ FC between early acute ZIKV infection and the convalescent stage was 6.135 and the average Log_2_ FC between late acute ZIKV infection and the convalescent phase was 1.378. Adjusted p-values for the upregulation of *CCL2* in early acute and late acute phases of the infection were 1.28E-65 and 0.00112, respectively. **(E)** Normalized counts of *HERC5* expression across each ZIKV infection time. The average Log_2_ FC was 3.14 between early acute ZIKV infection and convalescent period with an adjusted p-value of 5.87E-37. **(F)** Normalized counts of *CMPK2* expression across ZIKV infection times had an average Log_2_ FC of 3.3 between early acute and convalescent timepoints and an adjusted p-value of 2.42E-50.

We next compared gene expression between acute infection and the convalescent phase. Using a Log_2_ fold change (FC) with a cut-off value of 1, we identified 688 and 420 genes that were upregulated during early acute and late acute ZIKV infection, respectively (**Figure 1B**). Notably, 100 of these upregulated genes were common to both early and late acute ZIKV infection timepoints (**Figure 1B**). There were fewer genes downregulated at both early and late acute infection times when compared to the convalescent stage (**Figure 1C**). Specifically, we found that 166 and 276 genes were downregulated in early acute and late acute infection, respectively. There were 34 downregulated genes common to both early and late acute ZIKV infection (**Figure 1C**). Because we were interested in both shared and unique genes at each timepoint, we further characterize the infection stages by looking at the top differentially expressed genes.

We next combined all differentially expressed genes together and selected the top differentially expressed genes per infection time (supplementary tables with top 200 DEGs and a heatmap of the top 20 DGEs; **Table S3**, **Table S4**, **Figure S1**, and **Figure S2**). In examining genes with the highest upregulation as defined by Log_2_ FC, most genes were upregulated, and there was limited overlap between the two infection times. *CCL2* (Chemokine ligand 2) exhibited significant upregulation during the early acute stage with an average Log_2_ FC of 6.25 consistent with the immunological assays reported by Michlmayr *et al*., (**Figure 1D**) [46]. In late acute infection, *CCL2* was less upregulated but still statistically different with an average Log_2_ FC of 1.34 (**Figure 1D**). Additionally, top genes such as *HERC5*, *ATF3*, *CMPK2*, and *IFIT2*, displayed significant upregulation only during the early acute stage (**Figure 1E & 1F**, **Figure 3D**, **Figure S3**, and **Table S3**). Many of the top differentially expressed genes at the late acute time were unique to this stage of the infection. Of the top 20 genes in late acute stage only *IFI27* was in the top 100 genes during the early acute phase (**Figure S1 & S2**, **Tables S3 & S4**).

As shown in previous studies that investigated differential expression following viral infection, many of the top upregulated genes had roles in the innate immune response [47,48]. Notably, chemokines *CXCL10* and *CXCL11*, and interferon-stimulated genes *IFI27* and *IFIT1* were found to be significantly upregulated at both early and late acute times (**Figure 3E** and **Figure S3**). Overall, our differential gene expression analyses show unique expression patterns at early and late acute infection compared to the convalescent stage of ZIKV infection which reflect the earlier findings by Michlmayr et al. [46].

### Significant skipped exon alternative splicing events found in acute ZIKV infected pediatric PMBCs

While alternative splicing in cultured cells and human neural progenitor cells infected with ZIKV has been examined [47,52,66], the extent to which splicing alters the transcriptome in patients infected with ZIKV has not been extensively investigated. Using the RNA-Seq PBMC data [46] we examined the effects of ZIKV infection on alternative splicing in early and late acute patients relative to convalescent patients. Our analyses showed high levels of SE and mutually exclusive (MXE) alternative splicing events in both the early acute and late acute infection samples when compared to convalescent patients (**Figure 2A**). These data are consistent with our previous analysis showing that SE splicing was a major splicing event in ZIKV infected neuroblastoma SH-SY5Y cells [47]. Next, we examined the significant SE events (as determined by a FDR < 0.05 and ΔPSI > |0.1|) at the respective infection timepoints by principal component analysis and found a distinct clustering of the events in the early acute ZIKV infection samples while the late acute and convalescent had greater overlap (**Figure 2B**). Like the differential gene expression analyses, we observed that sex did not influence the SE splicing events in PBMC isolated from ZIKV infected patients (**Figure 2B**).

**Figure 2.**
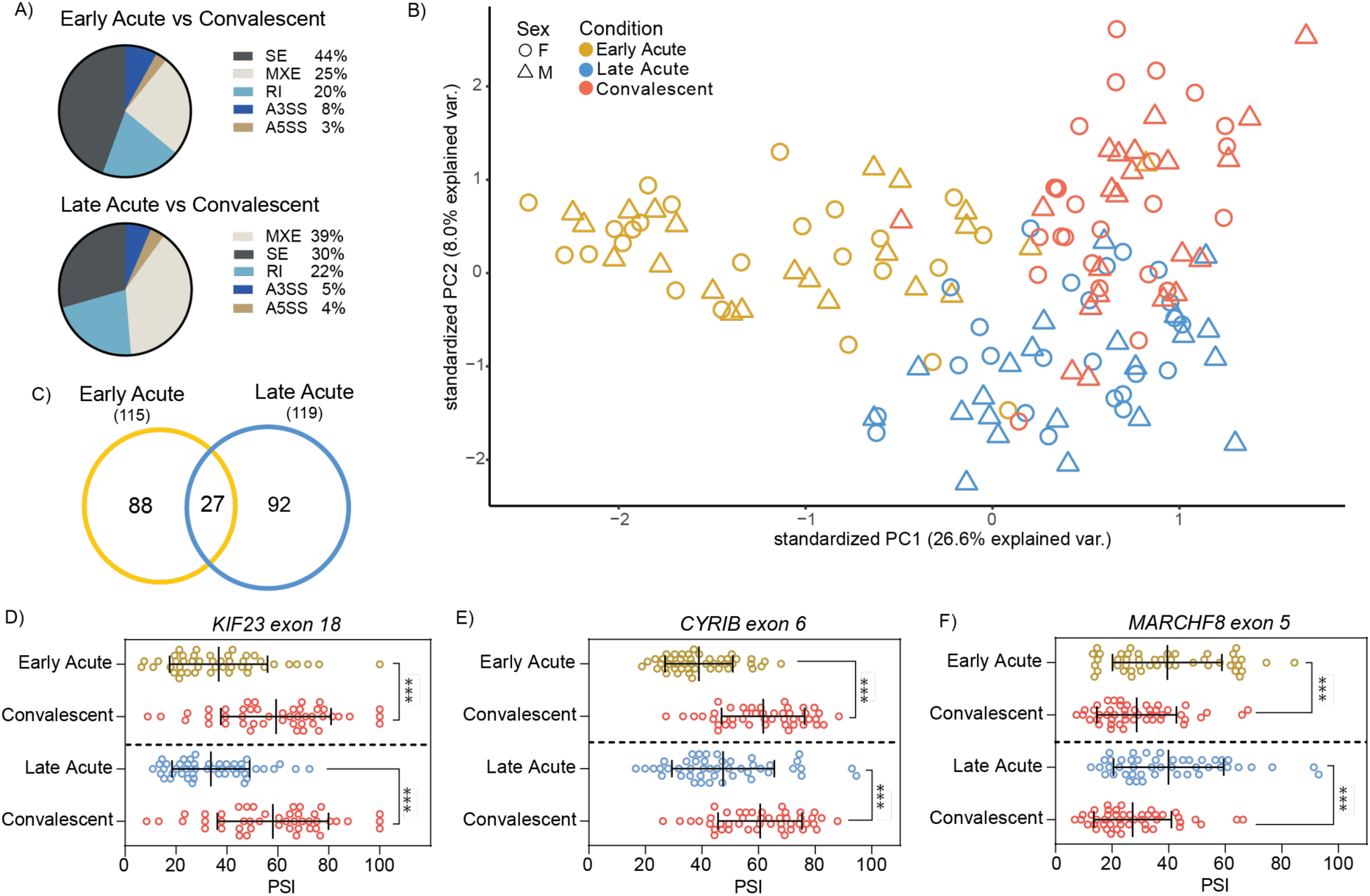
Alternative splicing analysis in ZIKV infected pediatric PMBC samples from early acute and late acute infection stages. **(A)** Pie charts show the percentage of each of the five alternative splicing events in the early acute and late acute patient samples when compared to patient samples from the convalescent phase. The alternative splicing events include skipped exons (SE), mutually exclusive exons (MXE), retained intron (RI), alternative to 3’ splice site (A3SS), and alternative to 5’ splice site (A5SS). **(B)** PCA plot PSI values from skipped exon events present between early acute vs convalescent, late acute vs convalescent, and early acute vs late acute timepoints. Samples also categorized by reported sex. **(C)** Venn diagram of overlapping significant skipped exon events (ΔPSI > |0.1| and <u>F</u>alse <u>D</u>iscovery <u>R</u>ate [FDR] < 0.05) between early acute and late acute infection when compared to convalescent. **(D-F)** Skipped exon splicing graphs with PSI values plotted for each patient sample at each infection timepoint are shown. The three genes chosen were **(D)** *KIF23* exon 18, **(E)** *CYRIB* exon 6, and **(F)** *MARCHF8* exon 5 and were selected based on being significantly alternatively spliced in both infection timepoints (Figure 2C). ***FDR < 0.05.

In examining the number of significant SE events relative to the convalescent phase, we found that the late acute phase of infection showed 92 distinct events compared to the 88 distinct events in the early acute phase, with 27 events overlapping between the early and late phase samples (**Figure 2C**). Within this overlap we observed three significant events from genes involved in regulation of the cytoskeleton and vesicular transport, which could influence virus assembly and egress and immune system processes [76–81]: kinesin family member 23 (*KIF23*), CYFIP related Rac1 interactor B (*CYRIB*), and membrane associated ring CH Finger 8 (*MARCHF8*). Exon 18 of *KIF23* and exon 6 of *CYRIB* were found to have significantly lower PSI values in early and late acute infections when compared to the convalescent stage (**Figure 2D-E**). Exon 5 of *MARCHF8* was found to have significantly higher inclusion in early and late acute when compared against convalescent (**Figure 2F**). These findings illustrate differences in SE alternate splicing events at different timepoints in ZIKV-infected patients which were previously unexplored by Michlmayr *et al*. [46].

### ZIKV infection of A549 cultured cells shares transcriptional profiles with patients during early and late acute stages of ZIKV infection

The molecular underpinnings of ZIKV are frequently studied in cultured cell lines. To determine if the transcriptomic events observed in ZIKV infected cell lines are similar to those in the pediatric patients we infected an A549 cell line with a contemporary ZIKV isolate namely PRVABC59 isolated from Puerto Rico [56] at a high multiplicity of infection and prepared samples for RNA-seq analysis 48 hours post-infection. Although A549 cells are derived from human lung adenocarcinoma, the cell line has an intact immune response and has been used for studies investigating the molecular virology and virus-host interactions of flaviviruses [82]. Moreover, infants with congenital Zika syndrome showed lung disease [83], and high viral titers were reported in the lung of mice infected with ZIKV [68,69]. Following RNA-seq, transcripts were mapped to the human genome and subjected to differential gene expression analysis.

A549 cells infected with ZIKV had 6,106 upregulated genes (Log_2_ FC > 2) and 2,027 downregulated genes (Log_2_ FC < 2). The notable upregulation of genes following ZIKV infection is consistent with the transcriptomic profile observed in the patient data (**Figure 1**) as well as in other cell model analyses [84]. When we compared the cell line and pediatric ZIKV infection RNA-seq data, we found 194 upregulated genes that were shared between at least two groups (**Figure 3A**). Early acute infection and A549 infected cells shared 150 upregulated genes including *ATF3* (**Figure 3D**), a stress induced transcription factor, as well as other genes involved in the innate immune response such as *MX1*, *DDX58*/*RIG-I*, *IFIT2*, *DHX58/MDA5*, *IFIT1* and *OASL* (**Supplementary Tables S3 & S5**). The overlap between patient samples collected during late acute phase and A549 infection included genes involved in response to immune stimulus such as *CCL16* and *IGHG1* (**Supplementary Tables S4 & S5**). Immune response genes such as the pro-inflammatory cytokine *CXCL10* (**Figure 3E**), *IFI27*, *USP18*, *IFIT1*, and *RSAD2/Viperin* were shared between both *in vivo* infection timepoints and A549 cells (**Figures S1 & S2**, and **Supplementary Tables S3-S5**). In Figure S3 we show the normalized counts and RT-qPCR validation of select genes shared between patient and A549 ZIKV infections. Overall, there was greater overlap in upregulated genes between the cell line and early acute patient infection.

**Figure 3.**
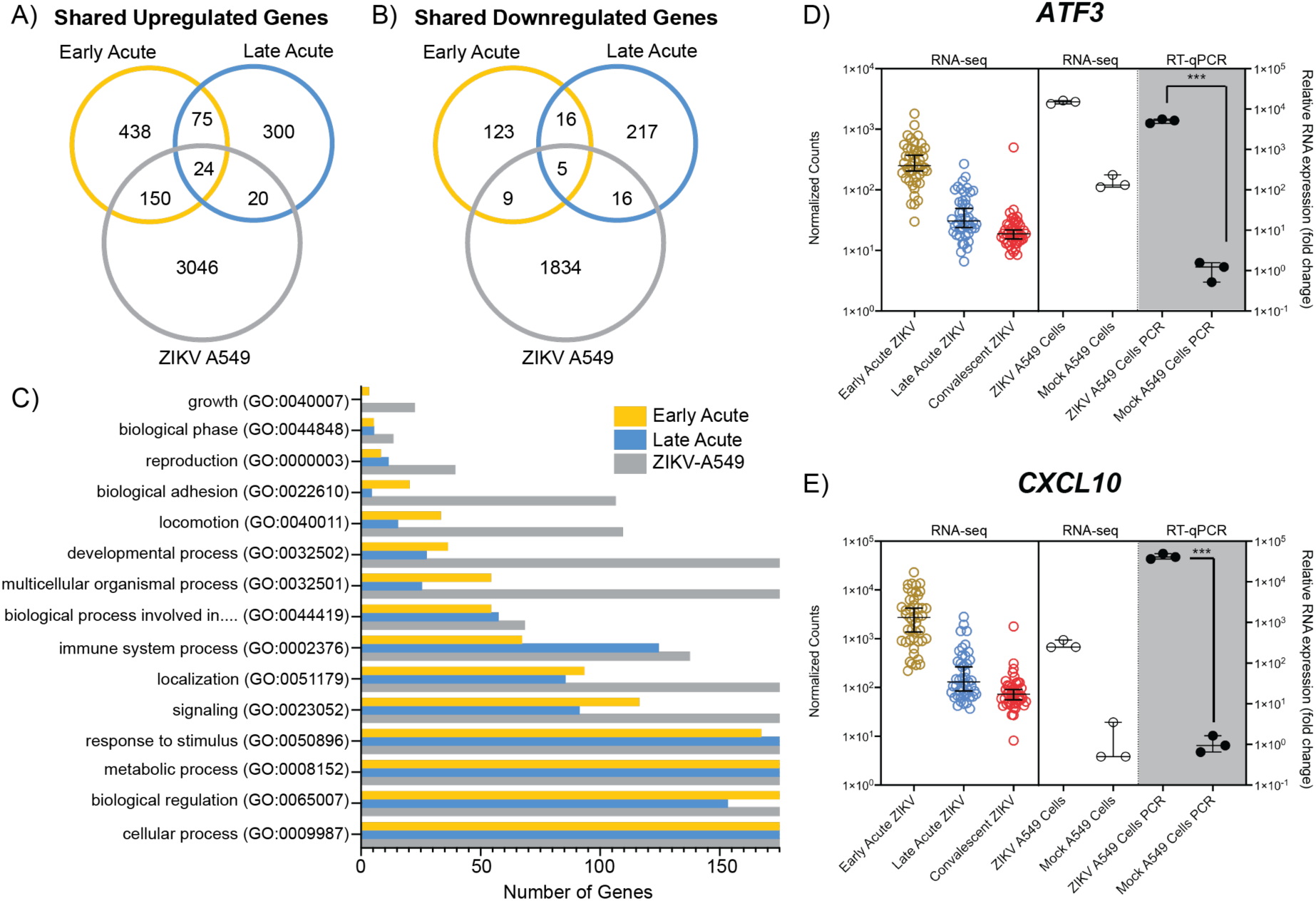
Analysis of differentially expressed genes following ZIKV infection in patients and an A549 cell model. **(A)** Venn diagram of shared genes upregulated following ZIKV infection in early acute and late acute patients (Log_2_ FC > 1) compared to ZIKV infected A549 cells (Log_2_ FC > 2). (**B)** Venn diagram of shared genes downregulated following ZIKV infection in early acute and late acute patients (Log_2_ FC < 1) compared to ZIKV infected A549 cells (Log_2_ FC < 2). **(C)** Gene Ontology (GO) biological process categories for differentially expressed genes from early acute, late acute and A549 ZIKV infections. Full category names and number of genes per category are found in Supplementary Table S6-S8. The GO biological process in (C) that has a truncated description represents “*biological process involved in interspecies interaction between organisms (GO:0044419)”*. **(D-E)** Normalized counts of **(D)** *ATF3* and **(E)** *CXCL10* expression across ZIKV infection times are shown in the left panel and the middle panel shows the data in ZIKV and mock infected A549 cells (middle panel). RT-qPCR validation of *ATF3* and *CXCL10* expression in A549 cells infected with ZIKV or uninfected (mock) is shown in the right shaded panel. Statistical significance of RT-qPCR was determined by student T-test. Error bars represent ± SD from three independent experiments. ***p<0.001.

We also compared the overlap in downregulated genes between patient infection timepoint and cell infection. Fourteen genes were shared between the early acute and A549 cell infections, and 21 genes between the late acute and cell line infections (**Figure 3B**). Of these, five genes were downregulated at both *in vivo* times and in the cell line infection. The five common genes included *SPP1*, *OPN*, as well as *ELFN2*, *RSPH14*, *CNTNAP3*, and *CNTNAP3B*. Overall, genes downregulated in ZIKV-infected A549 cells overlapped with both pediatric infection times, although we did not observe a clear preference for either early or late acute infection. We next examined gene ontology categories for the combined differentially expressed genes (upregulated and downregulated) from A549 cells, early acute and late acute ZIKV infections in patients. **Figure 3C** and **Supplementary Tables S6-S8** show the number of differentially expressed genes from the infection times and cell line for the top fifteen biological processes identified by Panther Gene Ontology [73]. Overall, the most abundant biological processes were conserved across the three groups and all three groups showed genes assigned to the top fifteen categories (**Figure 3C**). Notably, the GO term analyses identified cellular processes, biological regulation, metabolic process, and response to stimulus processes highly represented in differential gene expression from the patient samples and A549-infected cells. As expected, the robust ZIKV infection of A549 cells contributed a significantly higher number of genes in the respective fifteen GO categories (**Figure 3C**). In comparing the fifteen GO categories across the patient data, the infections in early acute patient samples showed a higher number of genes relative to the late acute patient samples (**Figure 3C**). Overall, the GO-categories show that ZIKV-infection in A549 cells modulates similar cellular processes as identified in early and late acute patient samples.

### ZIKV infected A549 cell line models alternative splicing changes found in infected patient data

Alternative splicing analysis was performed on the ZIKV infected A549 cells versus mock infection and compared to the splicing results of the early and late acute infection times in the pediatric PBMC data [46]. Most of alternative splicing events in the cell line detected were SE splicing events (67%, **Figure 4A , Figures S4-S6** and **Supplementary Tables S9-S11**) consistent with what was found when reanalyzing the patient RNA-seq data (**Figure 2A**). There were significantly more SE events in the ZIKV infected A549 cells versus either early acute or late acute infections in ZIKV infected patients (**Figure 4B**). This difference is likely due to the higher level of ZIKV infection in the cell line compared to ZIKV infection levels in the patients from the Michlmayr *et al* study [46].

**Figure 4.**
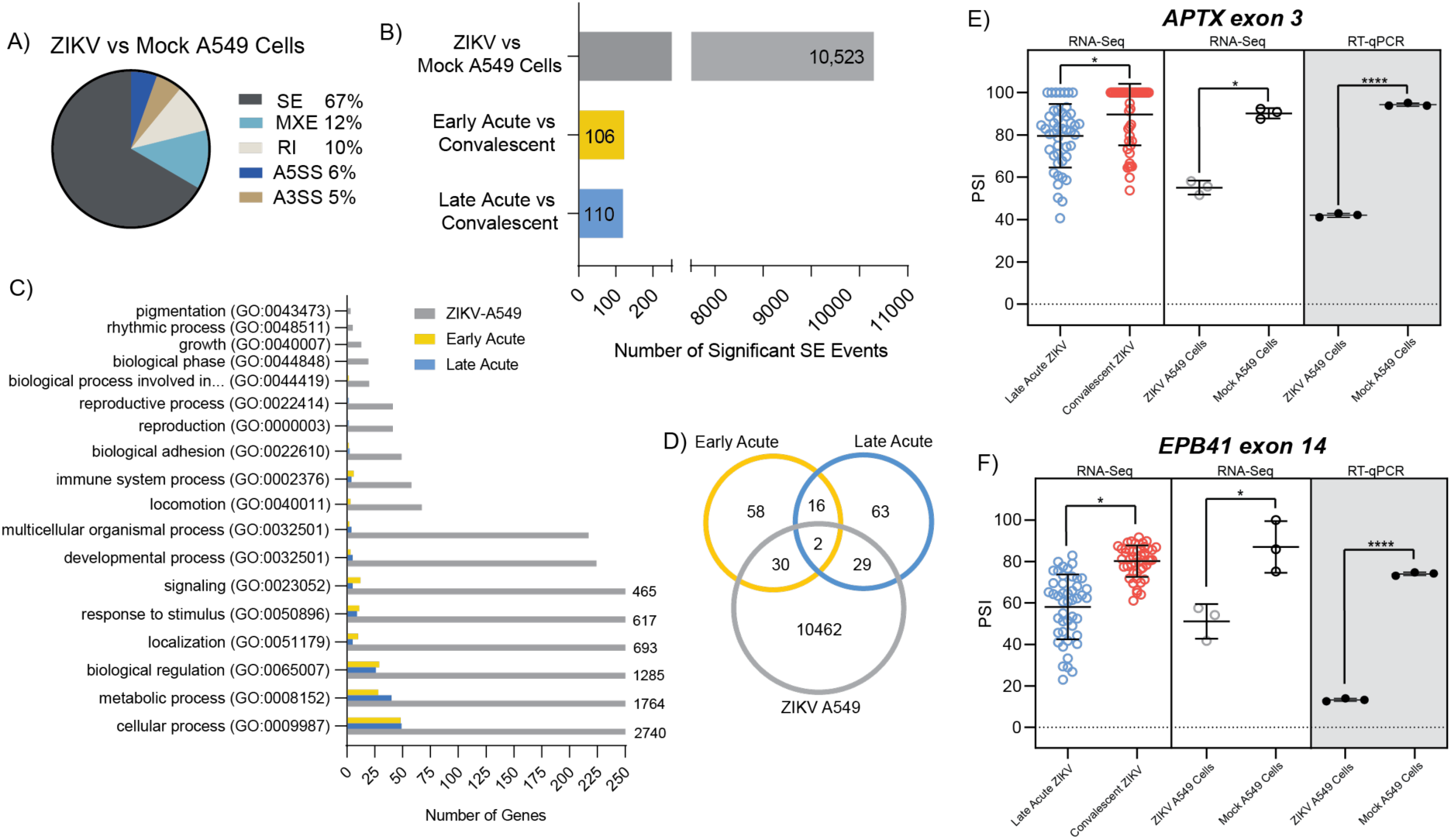
Comparison of alternative splicing changes between ZIKV infected patient samples and the A549 cell line infected with ZIKV. **(A)** Pie chart representing the percentage of each of the five alternative splicing events in the ZIKV infected A549 cells when compared to mock-infected A549 cells. (B) Bar chart showing the large number of significant skipped exon splicing events in the ZIKV-infected A549 cells and comparison to the two acute infection timepoints of the patient study relative to the convalescent samples. (C) GO pathways analysis of the significant skipped exon events for the ZIKV-infected A549 cells, the early acute, and the late acute infection timepoints ranked by number of genes found. The truncated fifth pathway is *biological process involved in interspecies interaction between organisms (GO:0044419)*. The number of genes within each GO term for the early and late acute genes are represented on the x-axis and the number of genes in ZIKV-infected A549 cells exceeding the x-axis are annotated on the chart. (D) Venn diagram showing the overlap of significant skipped exon events in the ZIKV-infected A549 cells and PBMCs from early and late acute infection timepoints in patients. (E-F) Splicing graphs for selected SE events namely Aprataxin *(APTX*) exon 3 and Erythrocyte Protein band 4.1 (*EPB41*) exon 14 with PSI values determined from the RNA-seq data reported by Michlmayr *et al* (left panel) [46]. PSI values from RNA-seq analysis of ZIKV infected A549 cells compared to mock (middle panel), and RT-PCR validation of each event in shaded portion of graph (right panel). ***FDR < 0.05. Significance of RT-PCR validation of the SE events was determined by three independent experiments and student T-test. ****p<0.0001.

Importantly, gene ontology enrichment analysis of the two datasets showed that the genes with SE events had shared pathways where cellular process, metabolic process and biological regulation were the notable GO term categories identified (**Figure 4C**). In examining the number of shared SE events between the groups, 37 SE events were shared between early and late acute phases, 41 events between early acute and A549-infected cells and 40 events between late acute and ZIKV-infected A549 cells. Of these, two SE events were shared in all data sets (**Figure 4D**). We attempted to validated by PCR these two specific spicing events, tryptophan tRNA synthetase 1 (*WARS1*) exon 1, and cyclin L1 (CCNL1) exon 6 (**Table S12**). Because the change in these two SE events occurred the ends of each transcripts, we were unable to design primers that would capture and validate these two splicing events. We were however able to show that exon 3 of aprataxin (*APTX*) and exon 14 of erythrocyte membrane protein 4.1 (*EPB41*) were excluded more frequently in late acute patient samples when compared to convalescent patient samples (**Figure 4E & 4F** and **Table S12**). We validated an additional four SE splicing events that were shared between ZIKV infected A549 cells and late acute patients namely *ATPSCKMT* exon 5, *FGFR1OP2* exon 5, *TBC1D14* exon 9, and *YTHDF1* exon 4 (**Figure S7**). Likewise, we show that exon 51 of *CEP290* and exon 9 of *RBSN* were significantly skipped in both ZIKV A549 infected cells and in early acute patients (**Figure S7** and **Supplementary Table S12**). Together these data show that infection in A549 cells may be used as an effective cell model to further study the molecular impact of alternative splicing on ZIKV infection.

## Discussion

In this study we re-examined the RNA-seq data from PBMCs collected from early acute, late acute, and convalescent ZIKV-infected patients [46] and compared these to RNA-seq data from A549 ZIKV infected cells. Our objectives were to investigate overlap between the data sets, and to determine if infection of cultured cell lines could be used to investigate transcriptional and alternative splicing changes in patients. Although our findings show that the number of genes differentially expressed in A549 infected cells was significantly greater, we found that the gene ontology pathways were similar in both cell and patient data (**Figure 3C** and **Figure 4C**). Moreover, A549 infected cells showed transcriptional changes that were reflected in both early and late acute infected ZIKV patients. We also examined alternative mRNA splicing profiles. Our data revealed that in early and late acute phases SE and MXE splicing events predominated in the patient samples (**Figure 2A**). In contrast only SE events were dominant in A549 infected cells (**Figure 4A**). However, our analyses did identify SE events common to both ZIKV-infected patients and cultured cells (**Figure 4**, **Figure S7** and **Supplementary Table S12**).

Following ZIKV infection, 688 and 420 genes were upregulated, and 166 and 275 genes were downregulated in early and late acute infected patients compared to convalescent samples, respectively (**Figure 1B & 1C**, and **Figure 3A & 3B**). Of interest, we did not observe any differences in gene expression when we considered sex as a variable (**Figure 1A**). The number of differentially expressed genes from patient PMBC samples were lower than the differentially expressed genes in ZIKV infected A549 cells (**Figure 3A & 3B**). The A549 cell line was robustly infected at a multiplicity of infection of 10 PFU/mL. Although the levels of ZIKV-infection in the patient PBMCs were not reported by Michlmayr *et al*., [46], a similar study of ZIKV infection in patient PBMCs showed a 3-4 log_10_ ZIKV RNA/mL 2-6-days post-infection [85]. Thus, the difference in the differentially expressed genes might in part be because of the difference in the virus load between the two systems. Despite the difference in total number of differentially expressed genes, genes associated with the immune response were altered in both systems. In Figure S3 we show normalized counts from the patient and cell derived RNA-seq data with subsequent RT-PCR validation of select immune response genes. These data are consistent with global transcriptome changes in other ZIKV RNA-seq data sets [47,50,51].

Of interest in the RNA-seq from the patients, we observed that in addition to CCL2 which was highlighted by Michlmayr and colleagues [46], we identified that interferon-induced transcripts *HERC5*, *CMPK2, ATF3* and *CXCL10* were also upregulated in ZIKV infected patients (**Figure 1D-1F** and **Figure 3D-3E**). *HERC5,* a critical enzyme involved in conjugating the interferon-induced ISG15 to different target proteins [86–88], is upregulated in ZIKV infected primary human brain microvasculature endothelial cells [89]. Interestingly, HERC5-directed ISG15 modification of ALIX and CHIMP4A, two proteins associated with the ESCRT pathway, resulted in the turnover of the two ESCRT proteins and increased tick-borne encephalitis flavivirus infection [90]. Cytidine/uridine monophosphate kinase 2 (CMPK2) is also a type I interferon stimulated gene that was previously shown to affect translation and restrict ZIKV infection [91–93]. Indeed, we also found in ZIKV infected A549 cells that CMPK2 was upregulated compared to mock infected cells (**Figure S3**). Similarly, *ATF3*, *CXCL10* and other innate immune response genes that were upregulated in ZIKV infected patients were found to be upregulated in A549 infected cells (**Figure 3D-3E**, and **Figure S3**). With such overlap, the cell culture system provides a useful platform to examine the molecular underpinning of such genes in ZIKV infection in cell culture. For example, Activating Transcription Factor 3 (ATF3) is a stress-activated transcription factor that is induced in response to a broad range of stress stimuli [94,95] and is upregulated in different ZIKV-infected cell types [47,66,89,96] To date however, the transcriptional control of ATF3 in response to viral infection is largely unknown. We recently determined that ATF3 is activated by ATF4, the master regulator of the integrated stress response, and functions to promote the expression of select immune response genes and restrict ZIKV infection [82]. Given that ATF3 is upregulated in patients during the early acute phase of ZIKV infection (**Figure 3D**), it is likely that ATF3 similarly contributes to the immune response in patients.

While transcriptional profiling in response to ZIKV infection in a plethora cell types and systems have been reported [47,48,50,51,97–99], only a few studies have examined the changes in alternative splicing of mRNAs in ZIKV infected cells [47,52,66]. Hu and colleagues previously examined the RNA-seq dataset from ZIKV infected human cortical neuroprogenitor cells (hNPCs) and determined that SE events followed by retained intron events comprised the highest type of alternative splicing event [52]. A similar distribution of alternative splicing events was also observed in ZIKV infected SH-SY5Y neuroblastoma and U87 glioblastoma cells [47,100]. Michlmayr and co-authors quantified the different isoforms of *CCL2* and report that four isoforms of CCL2 were differently express [46]. In particular, the *CCL2-201* isoform was shown to be elevated in early acute infection and then decreased in the late acute and convalescent stage of ZIKV infection. In our alternative splicing and skipped exon analyses we used the rMATS splicing program, rather than quantifying the isoforms of different transcripts. As a result of the difference in the analysis programs and log_2_(FPKM +1) values, we did not observe changes in the *CCL2* isoforms. In our analyses we determined that in both early and late acute ZIKV-infected patients compared to the convalescent stage, the highest type of alternative splicing event was SE, followed by MXE splicing and then retained intron events (**Figure 2A**). Consistent with other ZIKV infection of cultured cells [47,52,66], SE alternative splicing events were enriched in A549 infected cells (**Figure 4A**).

Towards our goal of establishing if ZIKV infection of cultured cells would mirror specific splicing events, we focused our analysis on the enriched SE events in both RNA-seq datasets from PBMC and A549 infected cells (**Figure 2A** and **Figure 4A**). We determined that seven skipped exon events were shared between early and late acute ZIKV infection, eight between early acute infection and A549 infected ZIKV cells, and fourteen between late acute infection and A549 ZIKV infected cells (**Figure 4D**). Indeed, for select transcripts that were shared between patient and A549 cell data sets we were able to validate the alternative splicing profile in A549 ZIKV infected cells (**Figure 4E-4F** and **Figure S7**). Curiously, an examination of these transcripts with our early analysis in SH-SY5Y cells and the recent report on alternative splicing in U87 infected cells revealed little overlap of SE events [47,66], suggesting that alternative splicing events in response to ZIKV might be cell type specific.

Interestingly, we identified two SE events, *WARS1* exon 1 and *CCNL1* exon 6, that were shared between patient and cell culture RNA-seq datasets (**Supplementary Table S12**). Due to the specific splicing events, we were unable to identify suitable primers to validate these events. However, from the RNA-seq analysis, the difference in the percent spliced in (ΔPSI) of exon 6 of Cyclin L1 (*CCNL1*) was -0.123 and -0.11 in early and late acute samples respectively, and -0.847 in A549 ZIKV-infected cells, indicating that exon 6 was likely spliced out during ZIKV infection. CCNL1 is thought to regulate RNA polymerase II transcription and influence splice site selection [101]. This protein encodes ten exons of which exon 6 is within the coding region. Thus, if exon 6 were excluded, protein expression and/or function would be affected to further perturb RNA splicing during flavivirus infection. Notably, this *CCNL1* skipped exon 6 event was not previously described in hNPC, SH-SY5Y and U87 ZIKV infected cells [47,52,66].

Tryptophan tRNA synthetase (WARS), catalyzes the addition of the tryptophan amino acid to the cognate tRNA [102]. There are two WARS paralogues that function in the cytoplasm (WARS1) and mitochondria (WARS2), respectively. Interestingly, early during pathogen infection secreted WARS1 was induced by interferon-γ and proposed to act prior to the complete initiation of the innate immune response pathway [103]. Similarly, WARS1 was shown to be secreted in response to vesicular stomatitis virus (VSV) infection to decrease virus protein levels and titers in immune cells [104]. Secreted full-length WARS1 was also reported as an entry factor for Enterovirus 71 [105]. Additionally, *WARS* was reported to be alternatively spliced in nasopharengeal swab samples collected from SARS-CoV-2 infected patients [106]. In our data we also found that *WARS1* was alternatively spliced in both early and late acute ZIKV infected patients, and in A549 infected cells (**Figure 4D** and **Table S12**). The ΔPSI of *WARS1* exon 1 in early acute, late acute and A549 ZIKV infected samples was 0.256, 0.113 and 0.557 respectively, indicating inclusion of this exon. Exon 1 encodes for sequences in the 5’ untranslated region and might harbor important elements that affect WARS1 protein expression. Despite the minimal overlap of alternatively spliced transcripts between three different ZIKV RNA-seq datasets, it is noteworthy that Brand and colleagues also reported the skipped exon 1 splicing event for *WARS* in Kunjin virus, Yellow Fever virus and ZIKV but not Sindbis virus-infected U87 cells [66]. *WARS* was similarly reported to be differentially spliced in primary neuronal cells infected with the highly virulent Tick-borne encephalitis flavivirus (TBEV) Hypr strain but not the Neudoerfl prototypical TBEV strain [107]. Since different aminoacyl tRNA synthetases have been reported to influence RNA virus infection [108], it is possible that *WARS* might have a more significant role to play during ZIKV infection than previously recognized. Last, it is noteworthy that select variations in *WARS* have been associated with microcephaly [109–111], a neurological defect associated with infants infected with ZIKV *in utero* [112–115], although it remains to be determined whether the alternative splicing of *WARS* exon 1 mirrors this phenotype.

In response to flavivirus infection the cellular transcriptome and alternative splicing landscape changes. Between different cell types the transcriptional changes particularly those relating to the immune response to ZIKV infection align [47,48,50,51,97–99], yet alternative splicing profiles show more variability [47,52,66]. Depending on the global or specific mRNA alternatively spliced profile, understanding of these different splicing events could broaden our knowledge of ZIKV pathogenesis. From studies with Dengue virus, the changes in alternative splicing are not merely in response to cellular immune or stress response, but rather are also directed by the RNA-dependent RNA polymerase NS5 that localizes to the nucleus and nuclear speckles [116,117]. Despite the knowledge that viral and cellular factors influence alternative splicing, there is presently a significant gap in understanding the consequence of differentially spliced mRNAs with respect to mRNA stability, translation, and localization in cell, all of which could have important effects on the virus infectious cycle and ZIKV pathogenesis.

## Acknowledgements

We thank The RNA Institute for the generous stipend support to GB as part of the RNA Institute Undergraduate Summer Research Fellowship Program. This research was supported by National Institute of Health grants to JAB (R01NS135254 and R01NS12048501) and CTP (R01 GM123050 and R21 AI178672). PB was supported by an America Heart Association Predoctoral Fellowship. The authors acknowledge Sawyer Hicks for advice on designing primers to validate alternative splicing events, and Drs. John Cleary and Marlene Belfort for the critical review, thoughtful comments, and feedback on the manuscript.

## Supplementary

**Figure S1. Differential gene expression of top 20 genes from early acute versus convalescent patient PBMCs.** Heatmap showing the top 20 differentially expressed genes between early acute and convalescent infection. Top genes were defined as smallest p-value with a log_2_fold change greater than 1.5. Yellow indicates high expression and darker blues indicate lower expression. The color bars on the top indicate timepoint with early acute as gold and convalescent in red. Log_2_ FC and p-values are provided in Table S3.

**Figure S2. Differential gene expression of top 20 genes from late acute versus convalescent patient PBMCs.** Heatmap showing the top 20 differentially expressed genes between late acute and convalescent infection. Top genes were defined as smallest p-value with a log_2_fold change greater than 1.5. Yellow indicates high expression and darker blues indicate lower expression. The color bars on the top indicate timepoint with late acute as blue and convalescent in red. Log_2_ FC and p-values are provided in Table S4.

**Figure S3:** Normalized gene counts for genes differentially expressed in PBMCs following ZIKV infection in patients and A549 cultured cells. Gene count plots for selected genes found upregulated in acute ZIKV infected patients and A549 cultured cells. Genes included are *CMPK2*, *CXCL11*, *DDX58/RIG-I*, *DHX58/MDA5*, *HERC5*, *IFIT, IFIT2*, *MX1* and *OASL*. The left panel of each graph includes normalized counts of acute ZIKV infected patients with points colored by infection time (gold – Early Acute, blue – Late Acute, and red – Convalescent). The right panel for each graph includes RNA-seq counts and RT-PCR validation in A549 cells. The y-axis on the left reports the gene count scale for patient and A549 ZIKV and mock infected cells. Average Log_2_ FC and p-values are provided in Tables S3-S5. The y-axis on the right reports the relative mRNA levels as a fold change relative to *ACTB*. The RT-qPCR data are from three independent experiments and error show ± SD. Statistical significance of RT-qPCR data was determined by student T-test. ***p<0.05

**Figure S4. Top 20 skipped exon events from early acute versus convalescent patient PBMCs.** Heatmap showing top 20 significant skipped exon events between early acute and convalescent infection. Significant events were defined as having a ΔPSI > 0.1 and FDR < 0.05 and top events were selected as the 20 greatest absolute ΔPSI values. Yellow indicates high percent spliced in values and dark blue indicates low percent spliced in values. The color bars on the top indicate timepoint with early acute as gold and convalescent in red. PSI and FDR values are provided in Supplemental Table S9.

**Figure S5. Top 20 skipped exon events from late acute versus convalescent patient PBMCs.** Heatmap showing top 20 significant skipped exon events between late acute and convalescent infection. Significant events were defined as having a ΔPSI > 0.1 and FDR < 0.05 and top events were selected as the 20 greatest absolute ΔPSI values. Yellow indicates high percent spliced in values and dark blue indicates low percent spliced in values. The color bars on the top indicate timepoint with early acute as gold and convalescent in red. PSI and FDR values are provided in Supplemental Table S10.

**Figure S6. Top 20 skipped exon events from ZIKV infected versus mock infected A549 cells.** Heatmap showing top 20 significant skipped exon events between ZIKV and mock infected cells. Significant events were defined as having a ΔPSI > 0.1 and FDR < 0.05 and top events were selected as the 20 greatest absolute ΔPSI values. Yellow indicates high percent spliced in values and dark blue indicates low percent spliced in values. The color bars on the top indicate timepoint with ZIKV infection as gray and mock infection in black. PSI and FDR values are provided in Supplemental Table S11.

**Figure S7. Validation of select skipped exon splicing events from ZIKV infected patients and A549 cultured cells.** Splicing graphs for selected events namely ATP synthase C subunit lysine N-methyltransferase *(ATPSCKMT)* exon 4, Centrosomal protein 290 *(CEP290)* exon 51, FGFR1 oncogene partner 2 *(FGFR1OP2)* exon 5, Rabenosyn, RAB effector *(RBSN)* exon 9, TBC1 domain family member 14 *(TBC1D14)* exon 9, and YTH N6-methyladenosine RNA binding Protein F1 (*YTHDF1*) exon 4 with percent spliced in (PSI) values determined from the RNA-seq data reported by Michlmayr *et al* (left panel) (47). PSI values from RNA-seq analysis of ZIKV-infected A549 cells compared to mock (middle panel), and RT-PCR validation of each event in shaded portion of graph (right panel). The data from A549 cells are from three independent experiments (n=3), and statistical significance of RT-PCR data were determined by student T-test. ***FDR < 0.05 and **p<0.05, ***p<0.001, ****p < 0.0001.

## Abbreviations

AS: Alternative Splicing
A3SS: Alternative to 3’ splice site
A5SS: Alternative to 5’ splice site
ΔPSI: Change in percent spliced in
FC: Fold change
FDR: False discovery rate
GO: Gene ontology
MOI: Multiplicity of infection
MXE: Mutually exclusive exons
PBMC: Peripheral blood mononuclear cells
PCA: Principal component analysis
SE: Skipped exon
RI: Retained intron
ZIKV: Zika virus

