## Supplementary Figures S1-S7 & Tables S1-S2 for "Zika virus infection in a cell culture model reflects the transcriptomic signatures in patients"

Figure S1. Differential gene expression of top 20 genes from early acute versus convalescent patient PBMCs.

Figure S2. Differential gene expression of top 20 genes from late acute versus convalescent patient PBMCs.

Figure S3: Normalized gene counts for genes differentially expressed in PBMCs following ZIKV infection in patients and A549 cultured cells.

Figure S4. Top 20 skipped exon events from early acute versus convalescent patient PBMCs.

Figure S5. Top 20 skipped exon events from late acute versus convalescent patient PBMCs.

Figure S6. Top 20 skipped exon events from mock- versus ZIKV-infected A549 cells.

Figure S7. Validation of select skipped exon splicing events from ZIKV-infected patients and A549 cultured cells.

### Tables

Table S1: Primers used for validation of differential gene expression by RT-qPCR analysis.

Table S2: Primers used for validation of alternative splicing by RT-PCR analysis.

Table S3: Top 200 DEGs of early acute infection relative to the convalescent phase. (TableS3_EA_ZIKV_top200.xlsx)

Table S4: Top 200 DEGs of late acute infection relative to the convalescent phase. (TableS4_LA_ZIKV_top200.xlsx)

Table S5: Top 200 DGE genes for ZIKV infected A549 cells relative to mock infected cells. (TableS5_A549_ZIKV_top200.xlsx)

Table S6: DGE gene ontology of early acute infection relative to the convalescent time point. (TableS6_DGE_GO_EA.xlsx)

Table S7: DGE gene ontology for late acute infection relative to the convalescent time point. (TableS7_DGE_GO_LA.xlsx)

Table S8: DGE gene ontology for ZIKV infected A549 cells relative to mock infected cells. (TableS8_DGE_GO_ZIKV_A549.xlsx)

Table S9: Top 20 Skipped Exon splicing events of early acute infection relative to the convalescent phase. (TableS9_EA_top20_SE_Events.xlsx)

Table S10: Top 20 Skipped Exon splicing events of early late infection relative to the convalescent phase. (TableS10_EA_top20_SE_Events.xlsx)

Table S11: Top 20 Skipped Exon splicing events for ZIKV infected A549 cells relative to mock infected cells. (TableS11_A549_top20_SE_Events.xlsx)

Table S12: Supplemental table for all significant SE events of interest based on overlap analyses. (TableS12_SignificantOverlap_SE_Events.xlsx)

Table S13: Alternative splicing gene ontology of early acute infection relative to the convalescent time point. (TableS13_AS_GO_EA.xlsx)

Table S14: Alternative splicing gene ontology of late acute infection relative to the convalescent time point. (TableS14_AS_GO_LA.xlsx)

Table S15: Alternative splicing gene ontology of ZIKV-infected A549 cells relative to mock-infected cells. (TableS15_AS_GO_ZIKV_A549.xlsx)

**Figure S1.**

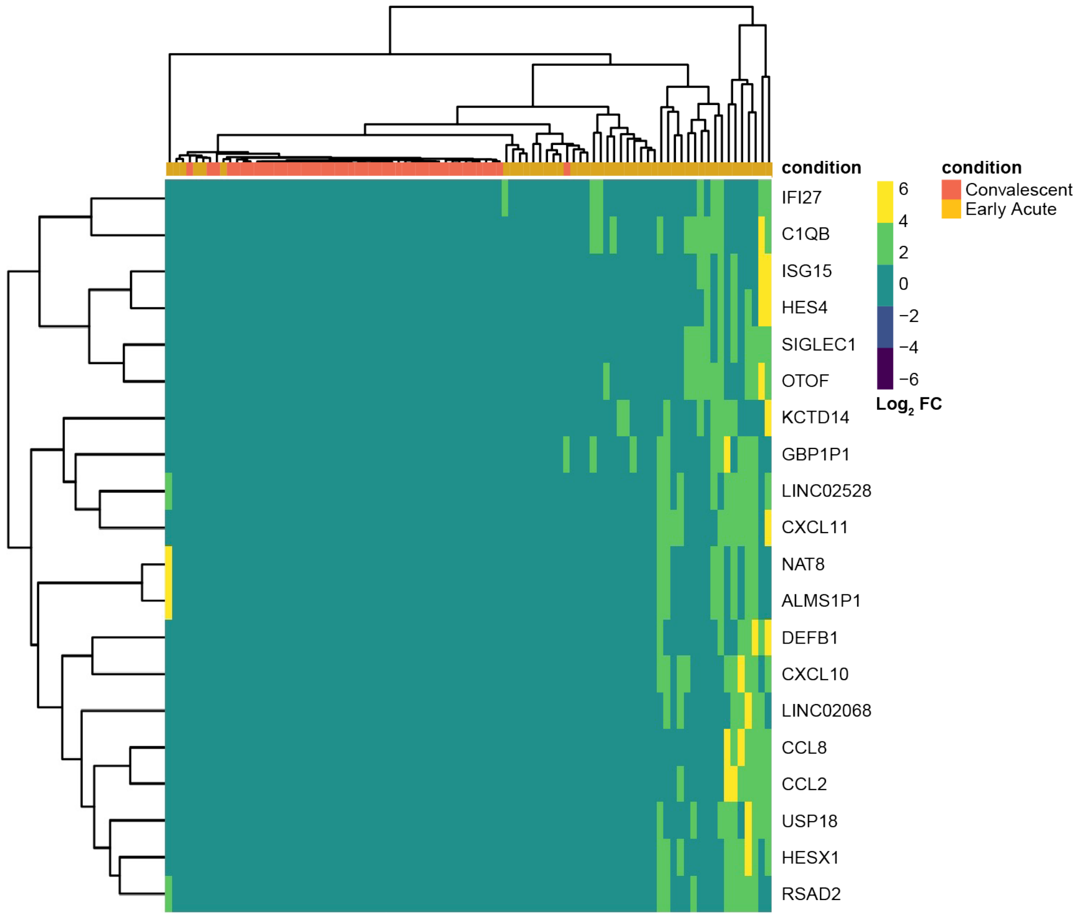


**Figure S1. Differential gene expression of top 20 genes from early acute versus convalescent patient PBMCs.** Heatmap showing the top 20 differentially expressed genes between early acute and convalescent infection. Top genes were defined as smallest p-value with a $\log_2$fold change greater than 1.5. Yellow indicates high expression and darker blues indicate lower expression. The color bars on the top indicate timepoint with early acute as gold and convalescent in red. $\text{Log}_2$ FC and p-values are provided in Table S3.

**Figure S2.**

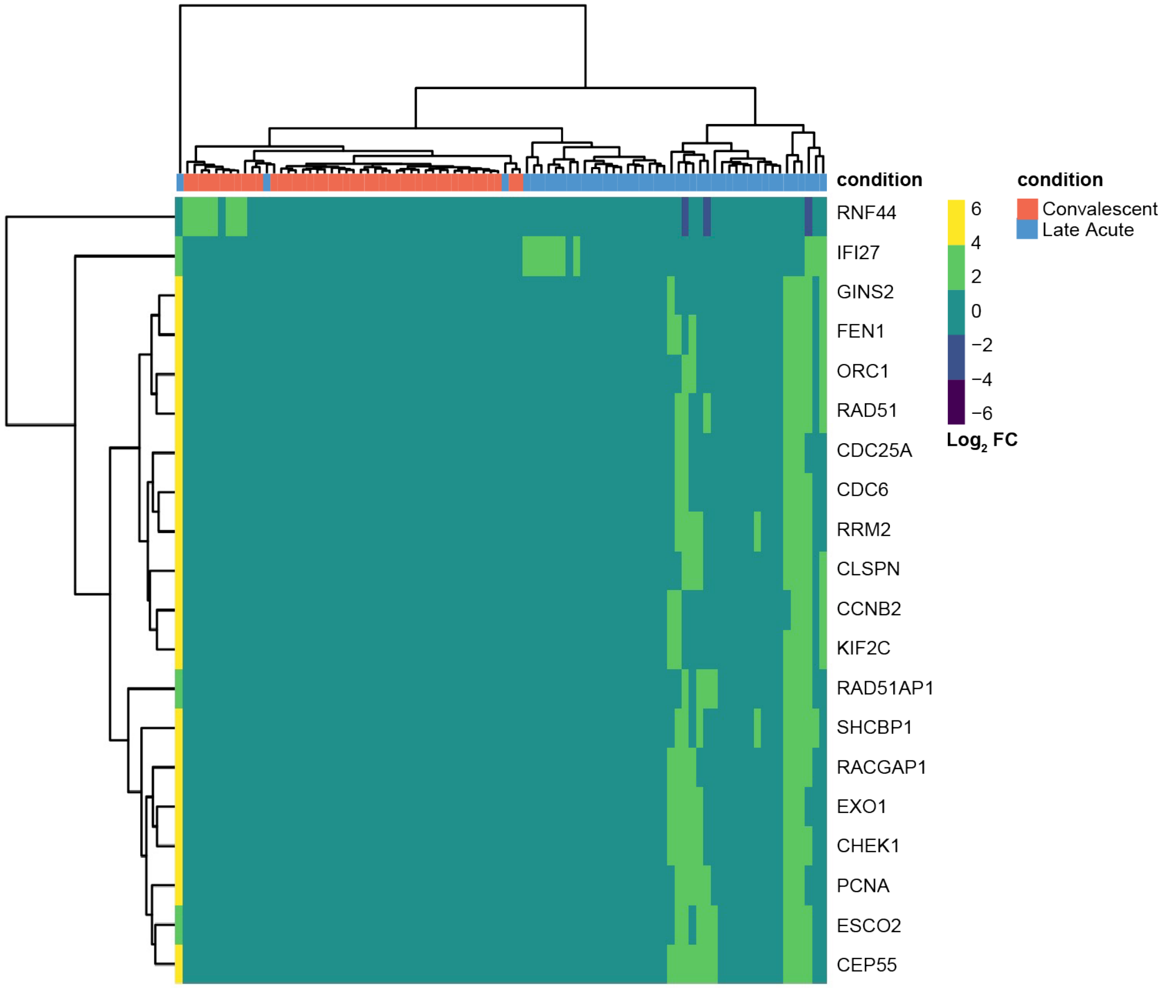


**Figure S2. Differential gene expression of top 20 genes from late acute versus convalescent patient PBMCs.** Heatmap showing the top 20 differentially expressed genes between late acute and convalescent infection. Top genes were defined as smallest p-value with a $\log_2$fold change greater than 1.5. Yellow indicates high expression and darker blues indicate lower expression. The color bars on the top indicate timepoint with late acute as blue and convalescent in red. $\text{Log}_2$ FC and p-values are provided in Table S4.

**Figure S3.**

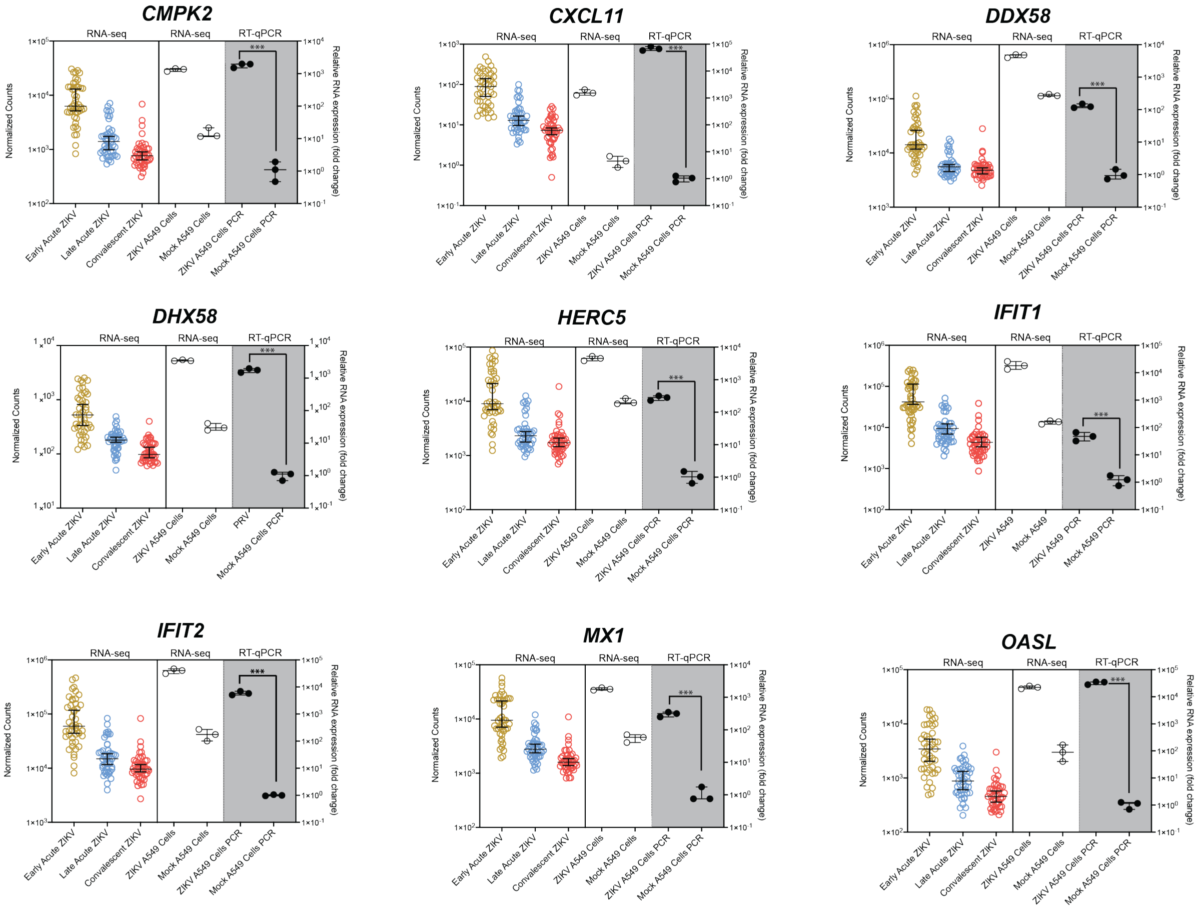


**Figure S3: Normalized gene counts for genes differentially expressed in PBMCs following ZIKV infection in patients and A549 cultured cells.** Gene count plots for selected genes found upregulated in acute ZIKV infected patients and A549 cultured cells. Genes included are *CMPK2*, *CXCL11*, *DDX58/RIG-I*, *DHX58/MDA5*, *HERC5*, *IFIT, IFIT2*, *MX1* and *OASL*. The left panel of each graph includes normalized counts of acute ZIKV infected patients with points colored by infection time (gold – Early Acute, blue – Late Acute, and red – Convalescent). The right panel for each graph includes RNA-seq counts and RT-PCR validation in A549 cells. The y-axis on the left reports the gene count scale for patient and A549 ZIKV and mock infected cells. Average $Log_2$ FC and p-values are provided in Tables S3-S5. The y-axis on the right reports the relative mRNA levels as a fold change relative to *ACTB*. The RT-qPCR data are from three independent experiments and error show ± SD. Statistical significance of RT-qPCR data was determined by student T-test. ***$p<0.05$

**Figure S4.**

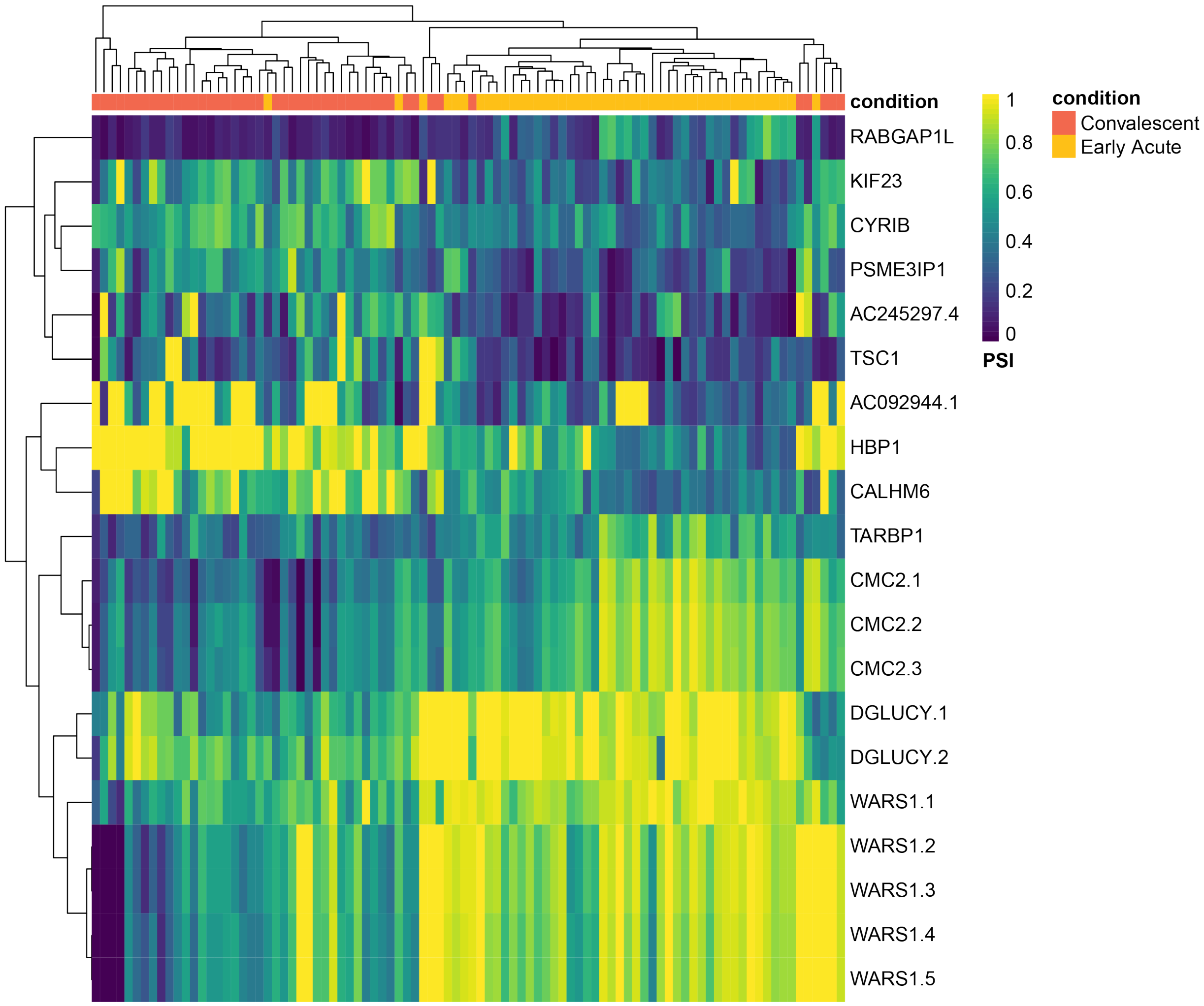


**Figure S4. Top 20 skipped exon events from early acute versus convalescent patient PBMCs.** Heatmap showing top 20 significant skipped exon events between early acute and convalescent infection. Significant events were defined as having a ΔPSI > 0.1 and FDR < 0.05 and top events were selected as the 20 greatest absolute ΔPSI values. Yellow indicates high percent spliced in values and dark blue indicates low percent spliced in values. The color bars on the top indicate timepoint with early acute as gold and convalescent in red. PSI and FDR values are provided in Supplemental Table S9.

**Figure S5.**

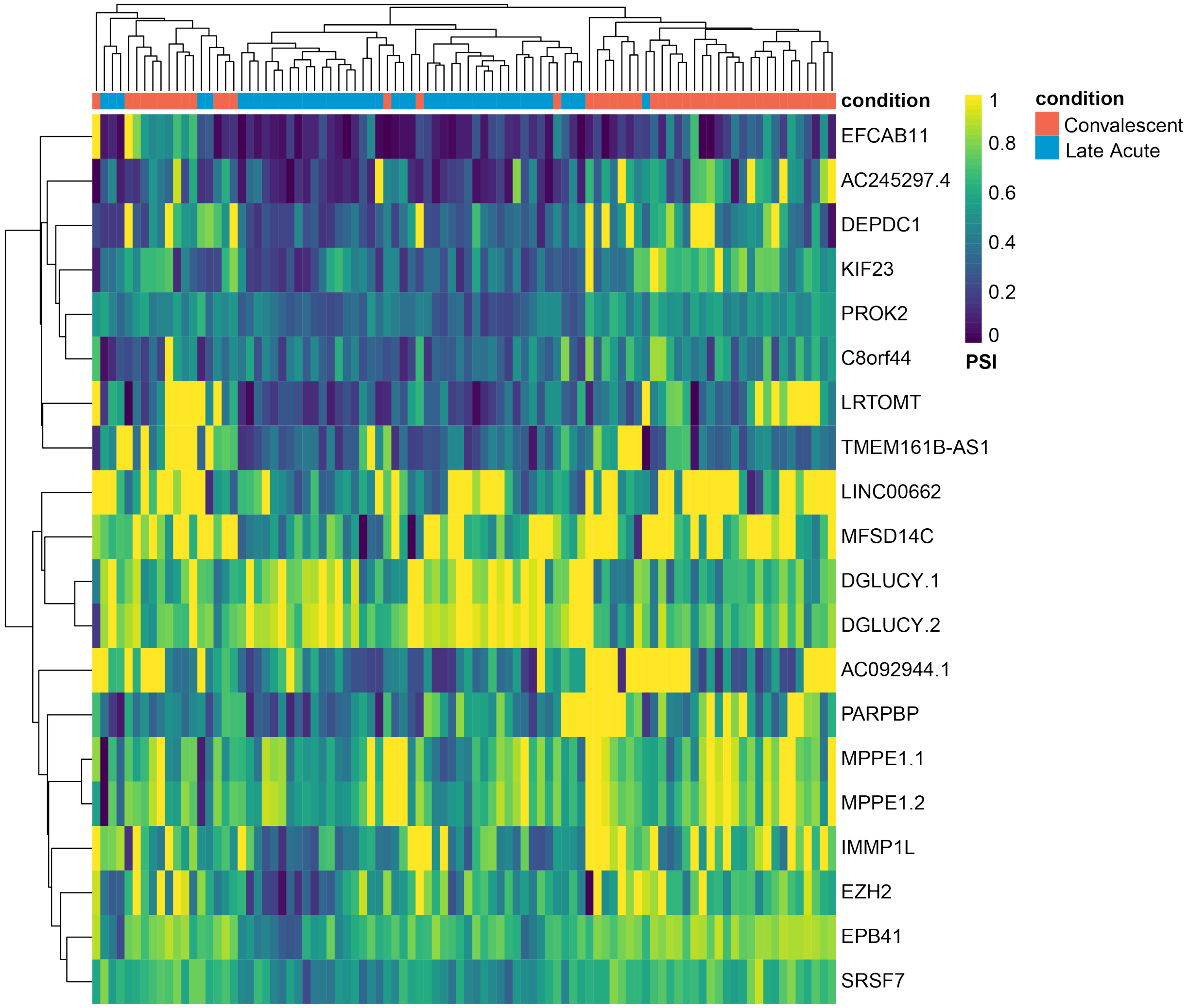


**Figure S5. Top 20 skipped exon events from late acute versus convalescent patient PBMCs.** Heatmap showing top 20 significant skipped exon events between late acute and convalescent infection. Significant events were defined as having a ΔPSI > 0.1 and FDR < 0.05 and top events were selected as the 20 greatest absolute ΔPSI values. Yellow indicates high percent spliced in values and dark blue indicates low percent spliced in values. The color bars on the top indicate timepoint with early acute as gold and convalescent in red. PSI and FDR values are provided in Supplemental Table S10.

**Figure S6.**

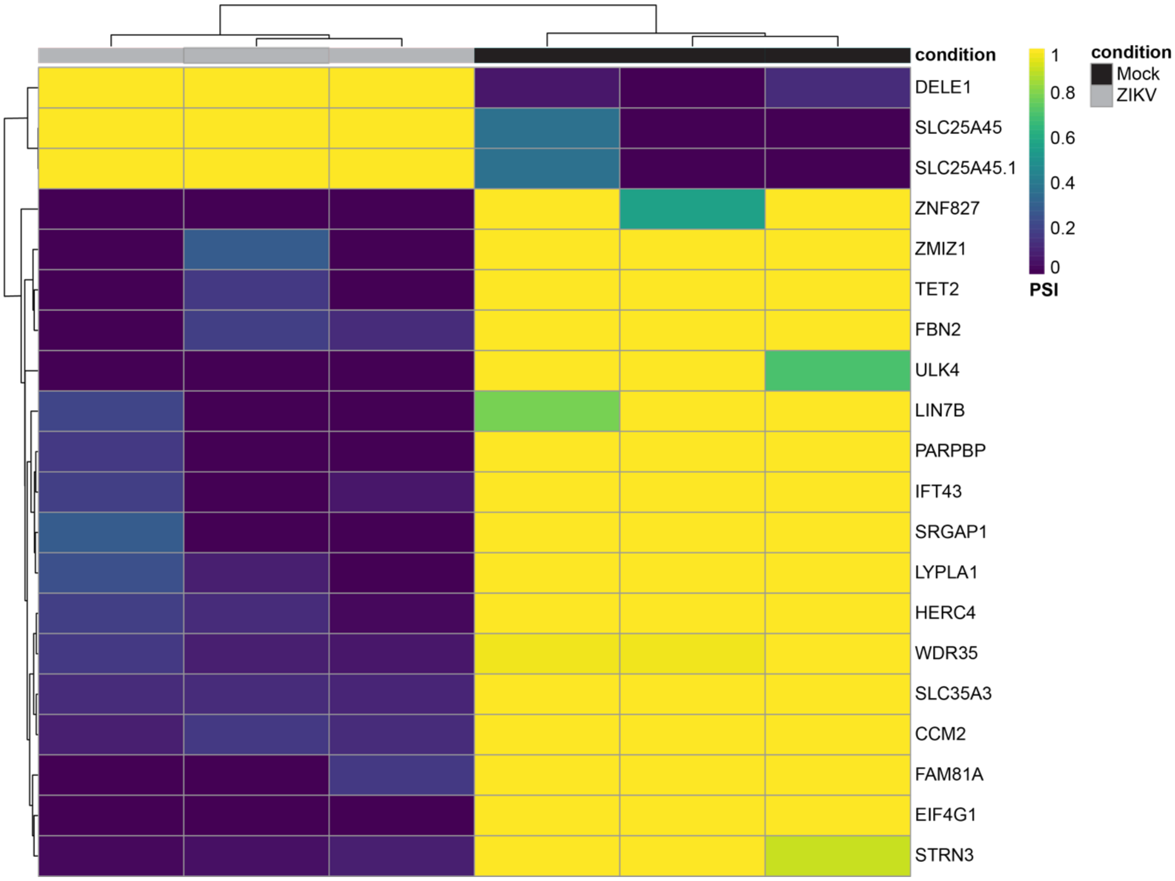


**Figure S6. Top 20 skipped exon events from ZIKV-infected versus mock-infected A549 cells.** Heatmap showing top 20 significant skipped exon events between ZIKV and mock infected cells. Significant events were defined as having a ΔPSI > 0.1 and FDR < 0.05 and top events were selected as the 20 greatest absolute ΔPSI values. Yellow indicates high percent spliced in values and dark blue indicates low percent spliced in values. The color bars on the top indicate timepoint with ZIKV infection as gray and mock infection in black. PSI and FDR values are provided in Supplemental Table S11.

**Figure S7.**

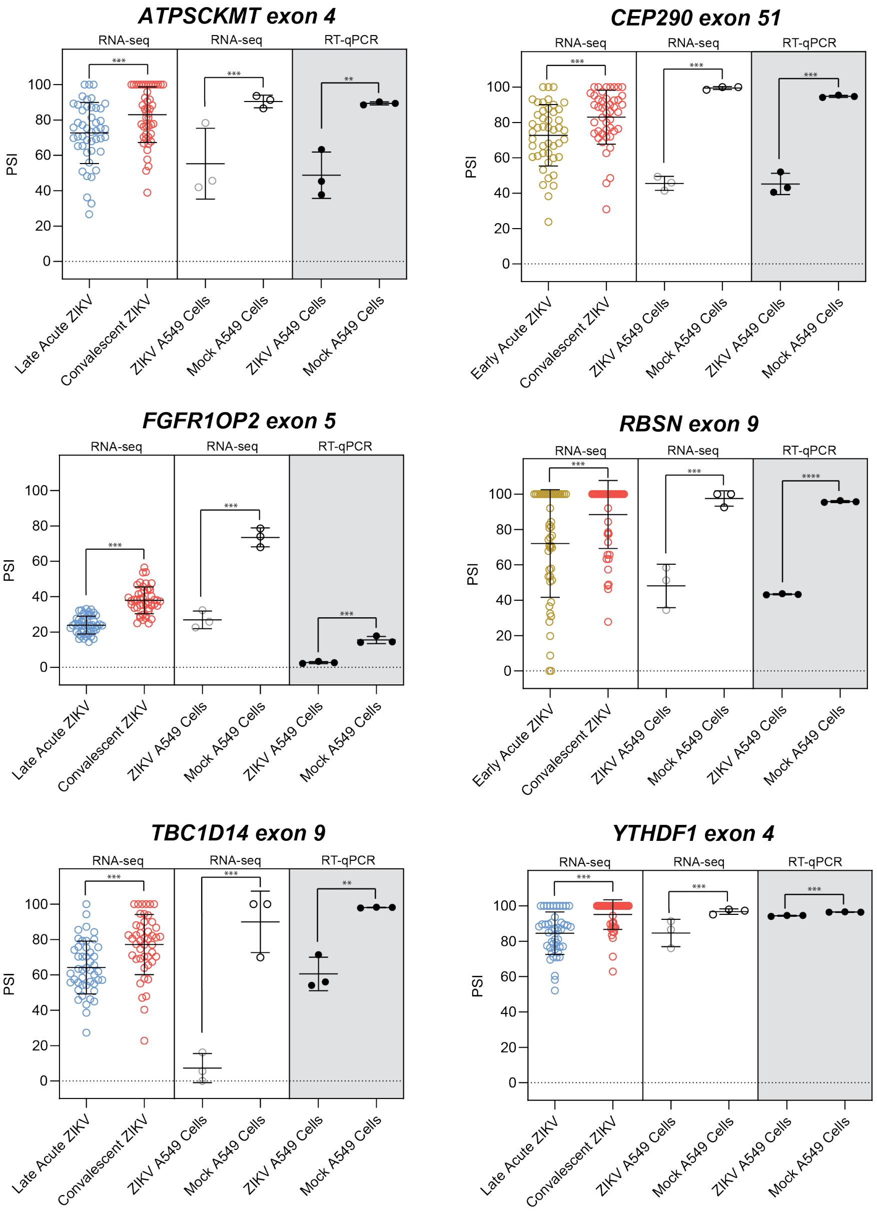


**Figure S7. Validation of select skipped exon splicing events from ZIKV infected patients and A549 cultured cells.** Splicing graphs for selected events namely ATP synthase C subunit lysine N-methyltransferase *(ATPSCKMT)* exon 4, Centrosomal protein 290 *(CEP290)* exon 51, FGFR1 oncogene partner 2 *(FGFR1OP2)* exon 5, Rabenosyn, RAB effector *(RBSN)* exon 9, TBC1 domain family member 14 *(TBC1D14)* exon 9, and YTH N6-methyladenosine RNA binding Protein F1 (*YTHDF1*) exon 4 with percent spliced in (PSI) values determined from the RNA-seq data reported by Michlmayr *et al* (left panel) (47). PSI values from RNA-seq analysis of ZIKV infected A549 cells compared to mock (middle panel), and RT-PCR validation of each event in shaded portion of graph (right panel). The data from A549 cells are from three independent experiments, and statistical significance of RT-PCR data were determined by student T-test. *FDR < 0.05 and **p<0.01, ***p<0.05, ****p < 0.0001. N=3.

**Table S1: Primers used for validation of differential gene expression by RT-qPCR analysis.**

| Gene name | Forward (5'-to-3') | Reverse (5'-to-3') |
|---|---|---|
| *Beta-actin (ACTB)* | GTCACCGGAGTCCATCACG | GACCCAGATCATGTTTGAGACC |
| *ATF3* | TGTCAAGGAAGAGCTGAGGTTTG | GATTCCAGCGCAGAGGACAT |
| *IFIT2* | AAGCACCTCAAAGGGCAAAAC | TCGGCCCATGTGATAGTAGAC |
| *MX1* | GGCATAACCAGAGTGGCTGT | CATTACTGGGGACCACCACC |
| *HERC5* | GACGAACTCTTGCACCGTCT | GCGTCCACAGTCATTTTCCAC |
| *DDX58* | AGAGCACTTGTGGACGCTTT | ATACACTTCTGTGCCGGGAGG |
| *CMPK2* | AGAAATGTCCTACCAGCGGA | AGGACCTTTTCTCTGGAGGGG |
| *OASL* | GCTGAAGGATGGGCAGAAATT | CACCCCCTGAGGTCTATGTGA |
| *CXCL10* | TGAATCCAGAATCGAAGGCCA | TGCATCGATTTTGCTCCCCT |
| *CXCL11* | GAAAGGTGGGTGAAAGGACCA | TGTTGGACTCCTTTGGGCAG |
| *DHX58* | AGGAACTGTGGGGAGGTCTG | CACGCGGGACCACTTTTTG |

**Table S2: Primers used for validation of alternative splicing by RT-PCR analysis.**

| Gene name | Exon | Forward (5'-to-3') | Reverse (5'-to-3') |
|---|---|---|---|
| APTX | 3 | TGTGCTGGTTGGTGAGACAG | TCACCTCTTGGTCCTTCCCA |
| EPB4 | 14 | TCAGGGTCAGGTTGCAGAAG | CATTCACTAGGCCGTGGTTCT |
| ATPSCKMT | 4 | AGGGTTCACAGCAGTTGGTT | AAGGGAACCGGCAAGCAATA |
| YTHDF | 4 | TGAGCCCTACCTTACTGGAC | ATGCTGAGAACGCAGGGTTT |
| FGFR10P2 | 5 | CATCAGTCGGCCTTGGAACT | CAACCCTGTTGCTCGTCAAT |
| TBC1D14 | 9 | TCGAGATTTATGGTGGCAGGG | TGCTGCTATGAAGGACATGC |
| CEP290 | 51 | TGCAGAAAGCAGAGGTCCA | GCTCCACTTTGGTCCTTGTTAG |
